# Genetic Instrumental Variable (GIV) regression: Explaining socioeconomic and health outcomes in non-experimental data

**DOI:** 10.1101/134197

**Authors:** Thomas A. DiPrete, Casper A.P. Burik, Philipp D. Koellinger

**Author notes:** The authors all contributed to the data analysis and writing of the paper. The authors contributed equally to this work.

## Abstract

Identifying causal effects in non-experimental data is an enduring challenge. One proposed solution that recently gained popularity is the idea to use genes as instrumental variables (i.e. Mendelian Randomization - MR). However, this approach is problematic because many variables of interest are genetically correlated, which implies the possibility that many genes could affect both the exposure and the outcome directly or via unobserved confounding factors. Thus, pleiotropic effects of genes are themselves a source of bias in non-experimental data that would also undermine the ability of MR to correct for endogeneity bias from non-genetic sources. Here, we propose an alternative approach, GIV regression, that provides estimates for the effect of an exposure on an outcome in the presence of pleiotropy. As a valuable byproduct, GIV regression also provides accurate estimates of the chip heritability of the outcome variable. GIV regression uses polygenic scores (PGS) for the outcome of interest which can be constructed from genome-wide association study (GWAS) results. By splitting the GWAS sample for the outcome into non-overlapping subsamples, we obtain multiple indicators of the outcome PGS that can be used as instruments for each other, and, in combination with other methods such as sibling fixed effects, can address endogeneity bias from both pleiotropy and the environment. In two empirical applications, we demonstrate that our approach produces reasonable estimates of the chip heritability of educational attainment (EA) and show that standard regression and MR provide upwardly biased estimates of the effect of body height on EA.

---

A major challenge in the social sciences and in epidemiology is the identification of causal effects in non-experimental data. In these disciplines, ethical and legal considerations along with practical constraints often preclude the use of experiments to randomize the assignment of observations between treatment and control groups or to carry out such experiments in samples that represent the relevant population [1]. Instead, many important questions are studied in field data which make it difficult to discern between causal effects and (spurious) correlations that are induced by unobserved factors [2]. Obviously, confusing correlation with causation is not only a conceptual error, it can also lead to ineffective or even harmful recommendations, treatments, and policies, as well as a significant waste of resources (e.g., as in [3]).

One important source of bias in field data stems from genetic effects: Twin studies [4] as well as methods based on molecular genetic data [5, 6] allow estimation of the proportion of variance in a trait that is due to linear genetic effects (so-called narrow-sense heritability). Using these and related methods, an overwhelming body of literature demonstrates that almost all important human characteristics, behaviors, and health outcomes are influenced both by genetic predisposition as well as environmental factors [7-9]. Most of these traits are “genetically complex”, which means that the observed heritability is due to the accumulation of effects from a very large number of genes that each have a small, often statistically insignificant, influence [10].

Furthermore, genes often influence several seemingly unrelated traits, a phenomenon called direct or vertical pleiotropy [11, 12]. For example, a mutation of a single gene that causes the disease phenylketonuria is responsible for mental retardation and also for abnormally light hair and skin color [13]. Pleiotropy is not restricted to diseases. All genes involved in healthy cell metabolism and cell division can be expected to directly influence a broad range of traits such as body height, cognitive ability and longevity, even if the effect on each of these traits may be tiny. Similarly, any gene involved in neurodevelopment and brain function is likely to contribute to human behavior and mental health in some way [14].

In addition to direct pleiotropic effects, genes can also have indirect or horizontal pleiotropic effects, where a genetic variant influences one trait, which in turn influences another trait [11]. The similarity of the genetic architecture of two traits is estimated by their genetic correlation (i.e. the correlation of the “true” effect sizes of all genetic variants on both traits) [15], which captures both direct and indirect pleiotropic effects [16-18]. Genetic correlations exist between many traits and often exceed their phenotypic correlations [19], giving rise to the concern that direct pleiotropy may substantially bias studies that do not control for genetic effects [20].

### Significance Statement

We propose Genetic Instrumental Variable (GIV) regression - a method that controls for pleiotropic effects of genes on two variables. GIV regression is broadly applicable to study outcomes for which polygenic scores from large-scale genome wide association studies are available. We explore the performance of GIV regression in the presence of pleiotropy across a range of scenarios and find that it yields more accurate estimates than alternative approaches such as ordinary least squares regression or Mendelian Randomization. When GIV regression is combined with proper controls for purely environmental sources of bias (e.g. using control variables and sibling fixed-effects), it improves our understanding of the causal relationships between genetically correlated variables.

If an experimental design is not possible, the gold standard in the presence of genetic confounds is to compare outcomes for monozygotic (MZ) twins [21, 22], who are by definition genetically (almost) identical [23]. In addition, this approach also controls for effects that arise from shared parental environment. However, the practical challenge is that such studies require very large sample sizes of MZ twin pairs because differences within MZ twin pairs tend to be small or non-existent. Furthermore, unobserved environmental differences between the twins or reverse causation can still lead to wrong conclusions in this study design.

Another popular strategy to isolate causal effects in non-experimental data is to use instrumental variables (IVs) [24]. Valid IVs are conceptually similar to natural experiments: They provide an exogenous “shock” on the exposure of interest to isolate the effect of that exposure on an outcome. Valid IVs need to satisfy two important conditions.^1^ First, they need to be correlated with the exposure conditional on the other control variables in the regression (i.e. IVs need to be “relevant“). Second, they need to be independent of the error term of the regression conditional on the other control variables and produce their correlation with the outcome solely through their effect on the exposure (the so-called exclusion restriction). In practice, finding valid IVs that satisfy both requirements is difficult. In particular, satisfying the exclusion restriction is challenging.

Epidemiologists have proposed to use genetic information to construct IVs and termed this approach Mendelian Randomization (MR) [25-28]. The idea is in principle appealing because genotypes are randomized in the production of gametes by the process of meiosis. Thus, conditional on the genotype of the parents, the genotype of the offspring are the result of a random draw. So if it would be known which genes affect the exposure, it may be possible to use them as IVs to identify the causal influence of the exposure on some outcome of interest. Yet, there are four challenges to this idea. First, we need to know which genes affect the exposure and isolate true genetic effects from environmental confounds that are correlated with ancestry. Second, if the exposure is a genetically complex trait, any gene by itself will only capture a very small part of the variance in the trait, which leads to the well-known problem of weak instruments [29, 30]. Third, genotypes are only randomly assigned *conditional* on the genotype of the parents. Unless it is possible to control for the genotype of the parents, the genotype of the offspring is *not* random and correlates with everything that the genotypes of the parents correlate with (e.g. parental environment, personality, and habits) [31]. Fourth, if direct pleiotropic effects of genes are the source of the confound, these genes could obviously not be used as IVs. One could try to isolate a subset of genes that only influence the exposure, but such attempts are still hindered by our limited knowledge of the function of most genes [27, 32, 33].

Recent advances in complex trait genetics make it possi-ble to address the first two challenges of MR. Array-based genotyping technologies have made the collection of genetic data fast and cheap. As a result, very large datasets are now available to study the genetic architecture of many human traits and a plethora of robust, replicable genetic associations has recently been reported in large-scale genome-wide association studies (GWAS) [34]. These results begin to shed light on the genetic architecture that is driving the heritability of traits such as body height [35], BMI [36], schizophrenia [37], Alzheimer’s disease [38], depression [39], or educational attainment (EA) [15].

High quality GWASs use several strategies to control for genetic structure in the population, and empirical evidence suggests that the vast majority of the reported genetic associations for many traits is not confounded by ancestry [40-43]. Polygenic scores (PGS) have become the favored tool for summarizing the genetic predispositions for genetically complex traits [15, 39, 44, 45]. PGS are linear indices that aggregate the estimated effects of all currently measured genetic variants (typically single nucleotide polymorphisms, a.k.a. SNPs). The effects of each SNP on an outcome are estimated in large-scale GWASs that exclude the prediction sample. Recent studies demonstrate that this approach yields PGS that begin to predict genetically complex outcomes such as height, BMI, schizophrenia, and EA [35-37, 39, 46]. Although PGS still capture substantially less of the variation in traits than suggested by their heritability [47] (an issue we return to below), PGS capture a much larger share of the variance of genetically complex traits than individual genetic markers. The third challenge to MR in the above list could in principle be addressed if the genotypes of the parents and the offspring are observed (e.g. in a large sample of parent-offspring trios) or by using large samples of siblings or dizygotic twins where the genetic differences between siblings are random draws from the parent’s genotypes. However, the fourth challenge (i.e. pleiotropy) remains a serious obstacle despite recent efforts to relax the exogeneity assumptions in MR [48-50].

Here, we address the implications of pleiotropy for modeling causal relationships using non-experimental data. We demonstrate that pleiotropy is a serious source of bias in ordinary least squares regression (OLS) and MR. We propose alternative estimation strategies that use PGS for the outcome of interest to reduce bias arising from pleiotropy. In particular, we propose a novel approach that we call Genetic Instrumental Variables (GIV) regression that can be implemented using widely available statistical software. GIV regression estimates practically useful upper and lower bounds for the causal effect of an exposure on an outcome even in the presence of substantial direct pleiotropy.

We begin by providing intuition and laying out the assumptions of our approach. We go on to show that GIV regression produces accurate estimates for the effect of the PGS on the outcome variable when the other covariates in the model are exogenous, when the true PGS is uncorrelated with the error term net of the included covariates, and when the GWAS sample sizes are sufficiently large relative to the number of SNPs. We then turn to the more complex case of when a regressor of interest (*T*) is potentially correlated with unobserved variables in the error term because of pleiotropy, and we show with evidence from a comprehensive set of simulations that the bias under these assumptions with GIV regression is gen-erally smaller than with OLS, MR, or what we will term an enhanced version of MR (EMR).

Next, we demonstrate the practical usefulness of our approach in empirical applications using the publicly available Health and Retirement Study [51]. First, we demonstrate that a consistent estimate of the so-called chip heritability [47] of EA can be obtained with our method. Then, we estimate the effects of body height on EA. As a “negative control,“ we check whether our method finds a causal effect of EA on body height (it should not).^2^

Formal derivations and technical details are contained in the Supporting Information (SI).

## Theory

### Intuition

To build intuition for our approach, we introduce the concept of the “true” PGS for *y* which would be constructed using the true effects of each SNP on *y*. In theory, the true SNP effects could be estimated in a GWAS on *y* in an infinitely large sample that is drawn from the same population as the prediction sample. The true PGS would capture the narrow-sense heritability of *y*. Of course, the true PGS is unknown. All one can practically obtain is a PGS from a finite GWAS sample that will capture a part, but not all, of the genetic influence on *y* because the effect of each SNP is estimated with noise. The attenuated predictive accuracy of practically available PGS [45, 47, 52] is conceptually similar to the well-known problem of measurement error in regression analysis. It has long been understood that multiple indicators can, under certain conditions, provide a strategy to correct regression estimates for attenuation from measurement error [53-55]. We show below that by splitting the GWAS sample into independent subsamples, one can obtain several PGS (i.e. multiple indicators) in the prediction sample. Each will have even lower predictive accuracy than the original score due to the smaller GWAS subsamples used in their construction, but these multiple indicators can be used as instrumental variables for each other, and the instruments will satisfy the assumptions of instrumental variable (IV) regression to the extent that the measurement errors (the difference between the true and calculated PGS) are uncorrelated. Standard two-stage least squares regression [24] (readily available in statistics software packages) using at least one valid IV for the PGS of *y* can then be employed to back out an unbiased estimate of the heritability of *y*.

Next, presume the matter of interest is not heritability, but the causal effect of some treatment *T* on *y*, where *T* is also heritable and some genes have direct pleiotropic effects on both. If these genes are not known and not controlled for, regressing j/onT would result in omitted variable bias.^3^ Suppose the effects of all genes that influence *y* through channels other than T would be known. Theoretically, one could estimate these effects in a GWAS on *y* that controls for *T* in an infinitely large sample. That information could be used to construct a “true conditional” PGS in a prediction sample. Adding the true conditional PGS to a regression of *y* on *T* in the prediction sample would effectively eliminate bias arising from direct pleiotropy. However, the true conditional PGS is also unknown and all we can practically obtain is a noisy proxy of it from a finite GWAS sample. While it is not guaranteed, the general conclusion of the literature is that the use of proxy variables is an improvement over omitting the variable being proxied [56, 57]. Furthermore, having a valid IV for the conditional score would potentially correct for its noise and get us closer to estimating the true causal effect of *T* on *y.* As before, a valid IV can be practically obtained by splitting the GWAS sample into independent parts and standard IV estimation techniques such as two-stage least squares can be used. We refer to this approach as conditional genetic IV regression (GIV-C).

If conditional GWAS results are not available, one can still add the unconditional PGS for *y* as a control variable and use IV regression with multiple indicators for this score to correct for measurement error. We refer to this as GIV-U (for unconditional). GIV-U still corrects for bias arising from direct pleiotropy, but this strategy will over control and result in estimates for *T* that are biased towards zero because the unconditional PGS also includes indirect pleiotropic effects of genes that affect *y* only because they affect *T.* Yet, extensive simulations show that the combination of GIV-C and GIV-U turn out to produce reasonable upper and lower bounds for the effect of *T* on *y* across a broad range of scenarios if the only sources of bias are pleiotropic genes.

The GIV strategy starts to break down when bias arises from unobserved non-genetic factors as well as from pleiotropic effects. We show below that both GIV and MR produce biased estimates in this case. Yet, we demonstrate that the combination of GIV-C and GIV-U still outperforms OLS and MR. Furthermore, the GIV approach has additional utility because it can be combined with other strategies to reduce the effects of environmental endogeneity (e.g., additional control variables or family fixed effects). We demonstrate that these combined strategies can potentially provide accurate information about the effects of an exposure in situations with both genetic and non-genetic sources of endogeneity. In contrast, the problems for MR that are produced by pleiotropy bias are not fixable in a similar manner.

### Assumptions

GIV regression builds on the standard identifying assumptions of IV regression [24]. In the context of our approach, this implies six specific conditions: (1) Polygenicity: The outcome is a genetically complex trait that is influenced by many genetic variants, each with a very small effect. (2) Complete genetic information: The available genetic data include all variants that influence the variable(s) of interest. (3) Genetic effects are linear: All genetic variants influence the variable(s) of interest via additive linear effects. Thus, there are no genetic interactions (i.e. epistasis) or dominant alleles. (4) Unbiased GWAS results: The available GWAS results are not systematically biased by omitted environmental variables that correlate with genetic ancestry. Failure to control for population structure can lead to spurious genetic associations [31]. (5) Non-overlapping samples: It is possible to divide GWAS samples into non-overlapping sub-samples drawn from the same population. (6) The genetic effects on *y* are the same in the GWAS and the prediction samples, i.e. the genetic correlation between samples is one.

### Estimating narrow-sense SNP heritability from polygenic scores

Under these assumptions, consistent estimates of the chip heritability of a trait^4^ can be obtained from polygenic scores (for full details, see SI section 2). If *y* is the outcome variable, *X* is a vector of exogenous control variables, and 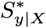 is a summary measure of genetic tendency for *y* in the presence of controls for *X*, then one can write

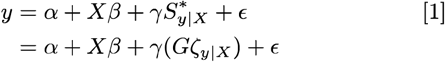

where *G* is an *n*×*m* matrix of genetic markers, and *ζ*_*y*|*X*_ is the *m*×1 vector of SNP effect sizes, where the number of SNPs is typically in the millions. If the true effects of each SNP on the outcome were known, the entire genetic tendency for *y* would be captured by the true unconditional score 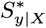, and the marginal *R*^2^ of 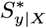 in equation 1 would be the chip heritability of the trait. We refer to the estimate of the PGS from actually available GWAS data in the presence of controls for *X* as *S*_*y*|*X*_ where

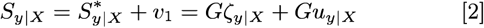

and *u*_*y*|*X*_ is the estimation error in *ζ*_*y*|*X*_ and *S*_*y*|*X*_ is substituted for 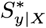 in equation 1. The variance of a trait that is captured by its available PGS increases with the available GWAS sample size to estimate *ζ*_*y*|*X*_ and converges to the SNP-based narrow-sense heritability of the trait at the limit if all relevant genetic markers were included in the GWAS and if the GWAS sample size were sufficiently large[45, 47, 52].

Equation 1 contains what is called in econometrics a “generated regressor” in that *S*\* is a function of a set of variables (*G*) and coefficients (*ζ*_*y*|*X*_) from another model. As previous work [58, 59] has established, OLS will provide consistent estimates of the parameters of equation 1 (though corrections for standard errors are needed) if *S* is substituted for *S*\* under a set of reasonable assumptions that include convergence in probability of 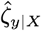 to *ζ*_*y*|*X*_ as the sample size grows larger. However, the practical utility of this mathematical result is questionable in the current context, when the number of variables in *G* are in the millions while the number of cases available to estimate *S*\* is far smaller than that. The imposed ratio of coefficients to cases requires non-conventional estimation methods that use a combination of statistical assumptions to obtain estimates of *S*. Empirical studies using PGS for a variety of traits have consistently demonstrated substantial attenuation in the estimate of *β* [45, 47, 52], and, while the bias diminishes with GWAS sample size, we are far away from sample sizes that bring this attenuation down to ignorable levels. This situation, therefore, calls for alternative strategies to address important questions with the datasets currently available.

The most straightforward solution to the problem of attenuation bias is to obtain multiple indicators of the PGS by splitting the GWAS discovery sample for *y* into two mutually exclusive subsamples with at least partially overlapping sets of SNPs. This produces noisier estimates of 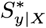 with lower predictive accuracy, but the multiple indicators can be used as IVs for each other. 2SLS regression using *S*_*y*1|*X*_ as an instrument for *S*_*y*2|*X*_ will then recover a consistent estimate of *β* in equation 1 under standard IV assumptions [55, 60].

As discussed more technically in the SI, an additional important assumptions for *S*_*y*1|*X*_ to be a valid instrument for *S*_*y*2|*X*_ is that *y* be a complex trait, meaning that it is influenced by a large number of genetic markers, each of which has a very small effect. If *y* is primarily influenced by a relatively small number of markers, then the method proposed here would not work well. However, there would also be no need for the proposed method, because the markers with large effects could be easily identified and their effects estimated with reasonable precision using discovery sample sizes that are already obtainable.

An additional required assumption is that the genetic markers are independent of each other. In general, genetic markers are correlated if they are located close to each other on the same chromosome. However, it is currently possible to isolate several hundred thousand markers from the total set of millions of SNPs that have sufficient spatial separation in the DNA to be essentially mutually independent, which means that this assumption can be satisfied to a sufficient level of accuracy.

Assuming that the variables in equation 1 are standardized to have mean zero and a standard deviation of one, and further assuming that the controls contained in *X* control for population stratification or are not correlate with genotype *G*, a consistent estimate of the chip heritability of *y* can now be obtained from 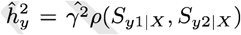, where *ρ* is the correlation coefficient. The heritability estimate 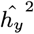 is not simply equal to 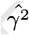 because we regressed on 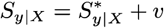. Thus, we standardize with the respect to the variance of 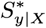 instead of 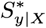 which leads to a bias equal to 1/*Var*(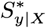). Multiplying 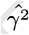 with the correlation between *S*_*y*1|*X*_ and *S*_*y*2|*X*_ recovers a consistent estimate for 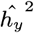 (see SI section 2.1).^5^

### Reducing bias arising from genetic correlation between exposure and outcome

Polygenic scores also play a potentially important role in situations where the question of interest is not the chip heritability of *y* per se, but rather the effect of some non-randomized exposure on *y* (e.g. a behavioral or environmental variable, or a non-randomized treatment due to policy or medical interventions). We can rewrite equation 1 by adding a treatment variable of interest *T*, such that

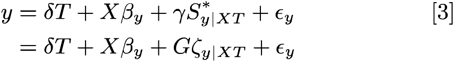

where

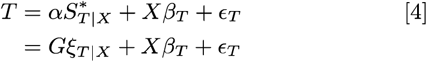

We assume that the disturbance term is uncorrelated with genetic variables.^6^ We now use the true conditional score 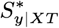 rather than 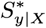 in the equation. Given that *T* is in the model, the effect of individual SNPs on *y* will generally involve a direct effect net of *T* (*ζ*) and an indirect effect stemming from the combination of their effect on *T* (*ξ*) and the effect of *T* on *y*. Having 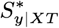 in the equation would effectively control for pleiotropic effects on *T* and *y*.

In standard MR, a measure of genetic tendency (*S*_*T*|*X*_) for a behavior of interest (*T* in equation 3) is used as an IV in an effort to purge 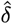 of bias that arises from correlation between *T* and unobservable variables in the disturbance term under the assumption that *S*_*T*|*X*_ is exogenous [60, 62]. One such example would be the use of a PGS for height as an instrument for height in a regression of EA on height. The problem with this approach is that the PGS for height will fail to satisfy the exclusion restriction if (some) of the genes affecting height also have a direct effect on EA (e.g. via healthy cell growth and metabolism) or if they are correlated with unmeasured environmental factors that affect EA.^7^

If the true conditional (net of *T*) genetic propensity for *y* could be directly controlled in the regression, pleiotropy would not bias MR coefficient estimates. For example, fixed effects regression where the same individual is observed multiple times would effectively control for pleiotropy (which does not vary over time), but this strategy is often not available (e.g., in a study of the effect of height on EA).^8^ Direct control for the conditional genetic propensity for *y* is, of course, not possible, because 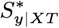 (more specifically, the coefficients, *ζ*_*y*|*XT*_ in equation 3) is not known. What is obtainable instead is a proxy for 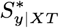 namely *ζ*_*y*|*XT*_ which contains measurement error due to finite GWAS sample size and potential bias in the estimate of *T* in the GWAS.

We refer to the combined use of *S*_*y*|*XT*_ as a control and *S*_*T*|*X*_ as an IV for *T* as “enhanced Mendelian Randomization” (EMR). However, controlling for *ζ*_*y*|*XT*_ as a proxy for 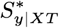 is not a perfect solution to pleitropy because it leaves a component of 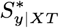 in the error term which is correlated with *S*_*T*_ due to pleiotropy.As a result, the EMR estimate for *δ* will be biased. The practical question, then, is whether alternative strategies that split the GWAS sample for *Y* to obtain multiple indicators of *S*_*y*|*XT*_ that can be used as IVs for each other (e.g. *S*_*y*_1_|*XT*_ and *S*_*y*_2_|*XT*_), are sufficient to rescue *S*_*T*_ as a practically useful IV for T. ^9^ This is a practical question beyond the reach of formal mathematics and best answered by simulation analyses. Unfortunately, and as we show in the SI, the pleiotropy-induced violation of the exclusion restriction when using genetic IVs for T is sufficient to produce serious bias in the estimated effect of *T* even if one attempts to control for pleiotropy using such strategies. The magnitude of bias clearly depends on the quality of the proxies for 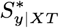. Yet, we find that pleiotropy leads to considerable bias in MR in virtually all scenarios we investigated (SI sections 2, 3, and 4, SI tables 2-15).

The problems posed by pleiotropy cannot be completely eliminated without knowledge of 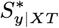 (SI section 2). However, this situation does not mean that the estimation problems introduced by pleiotropy are intractable. When endogeneity bias is driven by genetic correlation, the PGS for *y* can still be used to obtain more accurate estimates of the effect of *T* than can be obtained with MR, or, for that matter, with OLS that lacks controls for the direct effect of genetic markers on *y*. To gain insight into the best strategy, we consider the reasons for the pleiotropy bias. Regardless of whether the estimation strategy is OLS, or the second stage of IV regression involving OLS on predicted variables from the first stage, the coefficient bias comes from the extent to which the expected estimate of the OLS coefficients differ from the true coefficients,^10^ namely

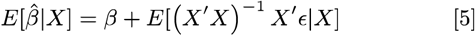

In other words, the coefficient bias from OLS is the expected regression coefficient of the error on the included variables in the regression. If *∊* is the sum of an omitted variable, *z*, which is correlated with the regressors, and additional variables that are uncorrelated with the regressors, then the bias for each coefficient *β*_*k*_ in equation 5 becomes the product of the regression coefficient for *x*_*k*_ in the regression of *z* on all the omitted variables multiplied by the effect of *z* on the outcome. For simplicity, we assume that the only variables in the regression are *T* and a potential proxy for 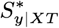, which we call 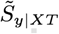. For any given proxy, 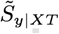, the bias in the estimate of 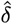 (the coefficient for *T* in equation 3) comes from the expected coefficient of *T* from a regression of 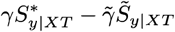 on *T* and 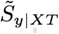. We consider three alternative approaches for the proxy 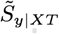, which we call simple OLS, GIV-C and GIV-U. First, we use *S*_*y*|*XT*_ as a proxy for 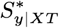 in a simple OLS regression. Second, we observe that *S*_*y*|*XT*_ is correlated with *T*; its inclusion in the error (by virtue of its being controlled) may affect the bias in 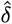 So we construct an estimate for *S*_*y1*|*XT*_ namely 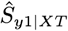, by using *S*_*y*2|*X*_(the unconditional PGS from the second GWAS sample) as its IV. We call this approach where 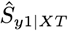 is used as the regressor in the second stage “conditional GIV regression” (GIV-C). We also employ a third estimator that uses the same IV as in GIV-C (i.e., *S*_*y*2|*X*_), but that substitutes the unconditional PGS for *y* (i.e., substitutes *S*_*y*1|*X*_ for the conditional PGS *S*_*y*1|*XT*_) in the structural model in equation 3. We then use *ζ*_*y*|*X*_ to predict *S*_*y*1|*X*_ obtaining 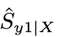 as the regressor in the second stage. We call this third approach “unconditional GIV regression” (GIV-U).

We generally expect the use of GIV-C to perform better than the use of the proxy *S*_*y*|*XT*_ in simple OLS. If the true effect of *T* on *y* is positive and positive pleiotropy is present, the estimated effect of *T* on *y* will have positive bias. This follows from the positive correlation between 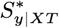 and *T* and from the positive effect of 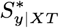 on *y*. The presence of the proxy *S*_*y*|*XT*_ in the first approach (simple OLS) adds a partially offsetting negative bias, because the correlation between *ζ*_*y*|*XT*_ and *T* is positive and the effect 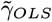 is also positive, but 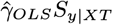 is being subtracted, which causes the offsetting bias to be negative. The net bias is expected to be positive, but we would expect it to be smaller with the inclusion of the proxy than with no proxy at all, both because the correlation between *T* and *S*_*y*|*XT*_ would be lower than between *T* and 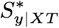 and because we expect 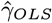 to be attenuated relative to *γ*. When GIV-C is used instead, the term in the error becomes 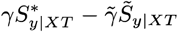 The presence in the first stage of GIV-C of *T*, which is correlated with 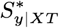 prevents the IV strategy from obtaining a consistent estimate of *γ*. Nonetheless, we would generally expect 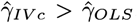 and therefore we expect the positive bias for the estimate of *δ* to be smaller when using GIV-C than when estimating *δ* using OLS and the proxy *S*_*y*1|*X*_. We confirm this in the simulations in the SI (SI Tables 2-7).

With GIV-U, the problem term in the error is 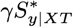 — 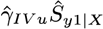 As before, the presence of the first term produces a positive bias in the estimate of *δ*, while the second term produces an offsetting negative bias. The offset will be stronger when the unconditional PGS for *y* is the regressor in the structural model, because the coefficients of the genetic markers in 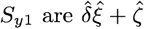, where *ζ* is the effect of the genetic marker on *T*. The presence of 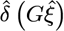 in the second endogenous term in the error (i.e., the second term in 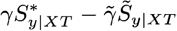) produces a stronger downward bias. This downward bias is made still stronger by the use of 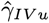 instead of 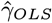 as the coefficient, because we expect the first stage regression to reduce the downward bias of 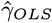. In other words, we expect these three proxies to behave differently in the simulations, and, as we will see, this expectation is met in practice. We establish via a comprehensive set of simulations that GIV-C and GIV-U provide upper and lower bounds for the effect of *T* across a range of plausible scenarios for pleiotropy and for heritability (SI Tables 2-7). We further establish through simulation analyses that GIV-C and GIV-U perform similarly in the case when endogeneity arises from pleiotropy and when it arises from pleiotropy in combination with genetic confounds for reasons other than pleiotropy (epistasis, effects from rare alleles, or genetic nurturing effects, where the environment of ego is shaped by genetically related individuals to ego [63]) (SI section 3, SI Tables 8-10).

## Simulations

We explored the performance of GIV regression in finite sample sizes using three sets of simulation scenarios (see SI appendix). The simulations generated genetic and phenotypic data at the individual level from a set of known models in a training sample and a hold-out sample using parameters that are realistic for genetically complex traits. We then estimated genetic effects on *T* and *y* using GWAS in the training sample and constructed polygenic scores with the estimated parameters for each SNP in the hold-out sample. Thus, the polygenic scores in our simulations have the realistic property that their predictive accuracy increases with the size of the training (i.e. GWAS) sample and the average effect size of each SNP [45, 52]. Finally, we analyzed the extent to which various estimation strategies recover the effect of the PGS for *y* on *y* and the effect of *T* on *y* in the hold-out sample. We produced these estimates using OLS, MR, EMR, proxy OLS, GIV-C and GIV-U regression, and we compared these results with the true answer across a range of parameter values. We ran 20 simulations with different random seeds for each set of parameters to obtain a distribution of estimated effects.^11^

The simulations specify that the true PGS scores for *y* and *T* covary as a result of genetic correlation. We made the conservative assumption that the entire genetic correlation between *y* and *T* is due to direct pleiotropy, i.e. all genes that are associated with both phenotypes have direct effects on both. In practice, this is unlikely to be the case, but it is equally unlikely that one can put a credible upper bound on (or completely rule out) direct pleiotropy.

In the first set of simulations, we assumed that the entire endogeneity problem arises from genetic confounds (SI section 2).

In the second set of simulations, we allowed endogeneity to arise from sources that are *both* correlated with genes and that cause the disturbance term in the structural equation for *y* to be correlated with the disturbance term in the structural model for *T* even if the true conditional PGS for *y* were included in the structural equation (SI section 3). This situation would occur if rare alleles were missing from the true PGS for *T* and the true conditional PGS for *y* based on known SNPs, and if the effect of these alleles was correlated with the true PGS scores for T and *y*. It would also occur under conditions of epistasis where nonlinear effects of genes were in the error and were correlated with the linear effects of genes in the PGS for *T* and the conditional PGS for *y*. Thirdly, this situation would occur if the genetic factors that affect T are correlated with the environmental factors in the disturbance term for *y* that are caused or selected by parental genes, which are correlated with the genes of sample members and therefore also with variables (like *T*) that are affected by the genes of sample members, i.e., by genetic nurturing [63].

In the third set of simulations, we specified the presence of a correlation between the error terms in the models for *y* and *T* that was not itself correlated with genetic variables (SI section 4). This would occur in a situation where some environmental or behavioral factor that is unrelated to genetics produces both an effect on *T* and an effect on *y*.

A summary of these results is in Table 1 for the case where the effect of *T* is set to 1.0 (See SI table 16 for details on the standardized effect size). Scenarios (A) to (D) refer to situations where pleiotropy is the only source of bias, scenario E contains pleiotropy plus other sources of genetic confounds, while scenarios (F) and (G) also include endogeneity from non-genetic (i.e. environmental) sources. The results provide considerable reason to be skeptical of estimates from MR. When pleiotropy is present, the MR strategy is undermined by the violation of the exclusion restriction for genetic IVs. Our results find that MR performs poorly even when non-genetic endogeneity is present along with pleiotropy. In contrast, GIV regression provides reasonable upper and lower bounds of the true effect of *T* on *y* if the source of endogeneity is only from pleiotropy or other genetic confounds (i.e. unobserved genetic variants, epistasis, or genetic nurturing) and the heritability of *T* and *y* is not extreme. GIV-C generally overestimates the effect of *T* but the overestimation is modest at low to moderate levels of pleiotropy and heritability and is more accurate than OLS without proxies, MR, or EMR. GIV-U generally underestimates the effect of of *T* on *y* but provides an estimate that is reasonably close to the true answer under conditions of low to moderate levels of pleiotropy and heritability. Even at higher levels of pleiotropy and heritability, the combination of GIV-C and GIV-U provides useful information about whether *T* actually has a causal effect on *y* and what the upper bound of this effect is likely to be.

**Table 1.** Illustrative Results from Simulations, Estimated coefficient for T.

|  | OLS | MR | GIV-C | GIV-U |
| --- | --- | --- | --- | --- |
| A Pleiotropy only<br>$h_y^2 = h_T^2 = 0.2$ | 1.1004<br>(0.0001) | 1.5040<br>(0.0012) | 1.0131<br>(0.0001) | 0.9419<br>(0.0001) |
| B Pleiotropy only<br>$h_y^2 = h_T^2 = 0.4$ | 1.2011<br>(0.0001) | 1.5024<br>(0.0004) | 1.0604<br>(0.0001) | 0.8575<br>(0.0001) |
| C Pleiotropy only<br>$h_y^2 = h_T^2 = 0.6$ | 1.3016<br>(0.0001) | 1.5020<br>(0.0002) | 1.1573<br>(0.0001) | 0.7263<br>(0.0002) |
| D Pleiotropy only<br>$h_y^2 = 0.8, h_T^2 = 0.2$ | 1.5004<br>(0.0005) | 3.4922<br>(0.0093) | 1.0776<br>(0.0002) | 0.8689<br>(0.0002) |
| E Genetic Confounds<br>$h_y^2 = h_T^2 = 0.5, \rho_{\nu y} = \rho_{\nu T} = 0.5$ | 1.3106<br>(0.0001) | 1.4609<br>(0.0002) | 1.1922<br>(0.0001) | 0.6422<br>(0.0002) |
| F Non-Genetic Confounds<br>$h_y^2 = h_T^2 = 0.5, \rho_e = 0.4$ | 1.4520<br>(0.0001) | 1.5032<br>(0.0002) | 1.4259<br>(0.0001) | 1.1193<br>(0.0001) |
| G Non-Genetic Confounds with control<br>$h_y^2 = h_T^2 = 0.5, \rho_e = 0.4, s = 0.5$ | 1.3643<br>(0.0001) | 1.5064<br>(0.0002) | 1.3346<br>(0.0001) | 0.9587<br>(0.0001) |

Mean estimated coefficient for T and standard error within parentheses of twenty simulations using different estimation methods for several models. For all models the genetic correlation ( $\rho$ ) was 0.5 and the coefficient for T ( $\delta$ ) was 1. $h_y^2$ and $h_T^2$ are the heritability parameters of y and T. $\rho_{\nu y}$ is the correlation between y and the genetic confound for y. $\rho_{\nu T}$ is the correlation between T and the genetic confound for T. $\rho_e = 0.4$ is the correlation between the non-genetic confound and y. $s$ is the share of the confound that is controlled for in terms of variance of the confound. These results are a selection from the SI tables, see the SI for all results. A: Table SI-2, B: Table SI-3, C: Table SI-4, D: Table SI-7, E: Table SI-9, F: Table SI-11 and E: Table SI-14. See row 2 of each table. See Table SI-16 for more details on the parameters, variance and standardized effect size.

In the case where the true value of *T* is zero (i.e., where the true model is equation 1), we expect that GIV-U will produce an estimate that is close to zero so long as the endogeneity comes either from pleiotropy or other genetic confounds (see SI for details). The simulations in the SI show that when the endogeneity is only from genetic sources, GIV-U estimates the effect of *T* to be close to zero regardless of the level of pleiotropy or inheritance that is specified in the simulations.

When the source of endogeneity is non-genetic in origin, we find that neither MR nor proxy controls for pleiotropy provide a satisfactory method for determining the effect of *T* on *y*. In this scenario, the pleiotropy creates endogeneity bias for genetic IV variables that defeats the ability of MR to solve the problem of non-genetic endogeneity via an IV strategy. Non-genetic endogeneity can cause even GIV-U to over-predict the effect of *T* on *y*, though in our simulations it is clearly the most accurate of all the estimators that we have surveyed when the effect of *T* is zero. Indeed, GIV-U always provides the most conservative estimate across the entire range of scenarios that we have surveyed, both for the case where an effect of *T* on *y* exists and when the effect of *T* is zero.

Inference with GIV can be further strengthened in cases where non-genetic endogeneity can be controlled either through observable variables or through strategies such as family fixed effects that reduce or eliminate the impact of non-genetic forms of environmental endogeneity. Indeed, environmental endogeneity is a concern in most applied research question that use non-experimental data. Reassuringly, our simulations show that GIV regression is a good estimation strategy in the presence of both direct pleiotropy and environmental endogeneity if control variables are available that manage to absorb a substantial share of the non-genetic confounds (SI tables 13-15). Therefore, we recommend using GIV regression always in combination with control variables the capture possible environmental confounds, ideally in datasets that allow controlling for family fixed-effects (e.g. using siblings or dizygotic twins). In the SI Appendix section 6, we provide additional practical guidelines for GIV regression.

The simulations described in the paper certainly do not cover all conceivable data generating processes, but they are nonetheless of considerable utility.

## Empirical applications

We illustrate the practical use of GIV regression in two empirical applications using data from the Health and Retirement Survey (HRS) for 2,751 unrelated individuals of North-West European descent who were born between 1935 and 1945 (SI appendix).

### The narrow-sense SIMP heritability of educational attainment

First, we demonstrate that GIV regression can recover the unbiased genomic-relatedness-matrix restricted maximum likelihood (GREML) estimate of the chip heritability of EA. Specifically, we follow the common practice in GREML estimates of heritability and analyze the residual of EA from a regression of EA on birth year, birth year squared, gender, and the first 10 principal components from the genetic data [64]. Next, we standardize the residual and regress it on a standardized PGS for EA using OLS or GIV. The results are displayed in Table 2. The OLS estimate of the PGS accounts for 6.8% of the variance in EA (*β*^2^=Δ*R*^2^=6.8%), which is substantially lower than the 17.3% (95% CI +/- 4%) estimate of chip heritability reported by [64] in the same data using GREML.^12^ Instead, the GIV regression results in columns 2 and 3 imply a chip heritability of 13.4% (CI +/- 3.9%) and 13.8% (+/- 4.0%), respectively. Thus, the 95% confidence intervals of the GREML estimate and the two GIV estimates overlap, demonstrating that GIV regression can recover the chip heritability of EA from polygenic scores.

**Table 2.**
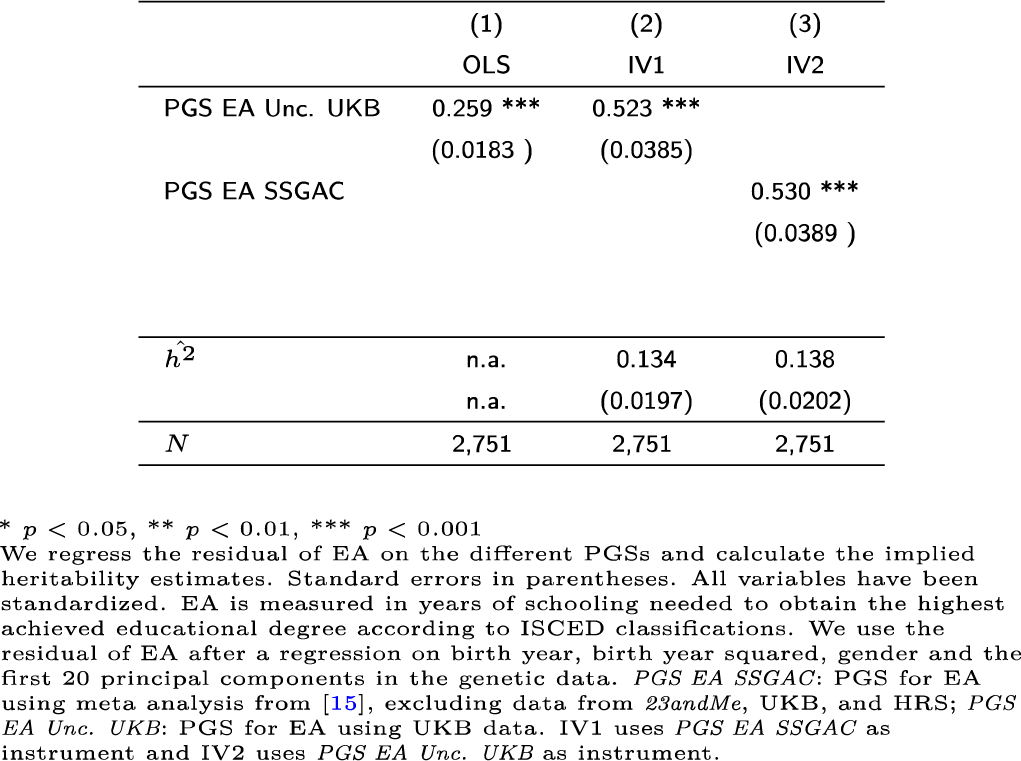
Effects of the PGS on (Residualized) Educational Attainment in the HRS subsample.

### The relationship between body height and educational attainment

Previous studies using both OLS and sibling or twin fixed effects methods have found that taller people generally have higher levels of EA [66-68]. They are also more likely to perform well in various other life domains, including earnings, higher marriage rates for men (though with higher probabilities of divorce), and higher fertility [69-74]. The question is what drives these results. Can they be attributed to genetic effects that jointly influence these outcomes? Are there social mechanisms that systematically favor taller or penalize shorter individuals? Or are there non-genetic factors (e.g., the uterine and post-birth environments especially related to nutrition or disease) that affect both height and these life course outcomes? The literature on the relationship between height and EA has found evidence that the association arises largely through the relationship between height and cognitive ability, which may suggest that the height-EA association is driven largely by genetic association between height and cognitive ability. We use GIV regression with individual-level data from the HRS to clarify the influence of height on EA, and we compare these results with those obtained from OLS and from MR. In addition, we conduct a “negative control” experiment that estimates the causal effect of EA on body height (which should be zero). A complete description of the materials and methods is available in the SI Appendix.

GWAS summary statistics for height were obtained from the Genetic Investigation of ANthropometric Traits (GIANT) consortium [35] and by running a GWAS on height (conditional on EA and unconditional on EA) in the UKB [75]. The UKB was not part of the GIANT sample. GWAS summary statistics for EA were obtained from the Social Science Genetic Association Consortium (SSGAC) for the unconditional PGSs. The most recent study of the SSGAC on EA used a meta-analysis of 64 cohorts for genetic discovery [15]. We obtained meta-analysis results from this study with the HRS, UKB and 23andMe cohorts excluded and we refer to the PGS constructed from these results as *EA SSGAC.* Furthermore, we obtained GWAS estimates for EA in the full UKB release (*N =* 442,183) from [76]. We refer to this PGS as *PGS EA Unc. UKB.* We also created a PGS for EA conditional on height by running a GWAS on EA in the same UKB sample (*PGS EA Cond. UKB).* There is sample overlap between *Height_GIANT* and *EA_SSGAC.* Therefore, whenever one of the two was used as regressor, we excluded the other as instrument and used a PGS from UK Biobank data instead to ensure independence of measurement errors in the PGS.

In Table 3 we report the estimated standardized effect of height on EA. The OLS results show that height appears to have a strong positive effect on EA, with 2.5 additional centimeters in height generating one additional month of schooling. MR appears to confirm the causal interpretation of the OLS result; indeed, the point estimate from MR is even slightly larger than from OLS. As discussed above, MR suf-fers from probable violations of the exclusion restriction due to pleiotropy. These violations could stem from the possibility that some genes have direct effects on both height and EA.1 They could also stem from the possibility that the PGS for height by itself is correlated with the genetic tendency for parents to have higher EA and income, which enables higher parental investments into their children who may therefore be more likely to reach their full cognitive potential and have higher EA. Controlling for the PGS is an imperfect strategy for eliminating this source of endogencity because the bias in the estimated effect of the PGS score also biases the estimated effect of height.

**Table 3.** Estimates of the Effect of Height on EA.

|  | (1)<br>OLS | (2)<br>MR | (3)<br>GIV-C | (4)<br>GIV-U |
| --- | --- | --- | --- | --- |
| Height | 0.136***<br>(0.0262) | 0.160***<br>(0.0481) | 0.168***<br>(0.0264) | 0.110***<br>(0.0262) |
| PGS EA Cond. UKB |  |  | 0.396***<br>(0.0367) |  |
| PGS EA Uncond. UKB |  |  |  | 0.384***<br>(0.0354) |
| $N$ | 2,751 | 2,751 | 2,751 | 2,751 |

\* $p < 0.05$ , \*\* $p < 0.01$ , \*\*\* $p < 0.001$
Standardized effect sizes and standard errors in parentheses. Birth year, birth year squared, gender, EA mother, EA father and the first 20 principal components are included as control variables. For MR, a PGS for height from UKB data was used as instrument for height. For GIV-C and GIV-U *PGS EA SSGAC* was used as an instrument.

If all of the bias in EA came from positive pleiotropy, then we would expect GIV-C and GIV-U to provide upper and lower bounds for the true effect of height on EA, respectively. Thus, if the only source of endogeneity is pleiotropy, the results in Table 3 would suggest that the truestandardized effect is between 0.11 (from GIV-U) and 0.17 (from GIV-C).

However, the negative control regression results in Table 4 provide substantial evidence of endogeneity bias from environmental sources. Given that the true effect of EA on height should be zero, then GIV-U would accurately estimate this effect to be zero in the absence of environmental endogeneity. Instead, GIV-U is reporting a significant positive effect of EA on height. MR is also reporting a positive and statistically significant effect of EA on height. This upward bias in the MR estimate is strong evidence of pleiotropy bias that invalidates the IV in MR. The guidance from the simulations points to a true estimate of the effect of height on EA that is as small or smaller than the GIV-U estimate, which is 25% smaller than the estimate from MR. The extent of upward bias in the GIV-U estimate depends on the strength of environmental variables that simultaneously affected the height of HRS respondents and also affected their EA.

**Table 4.**
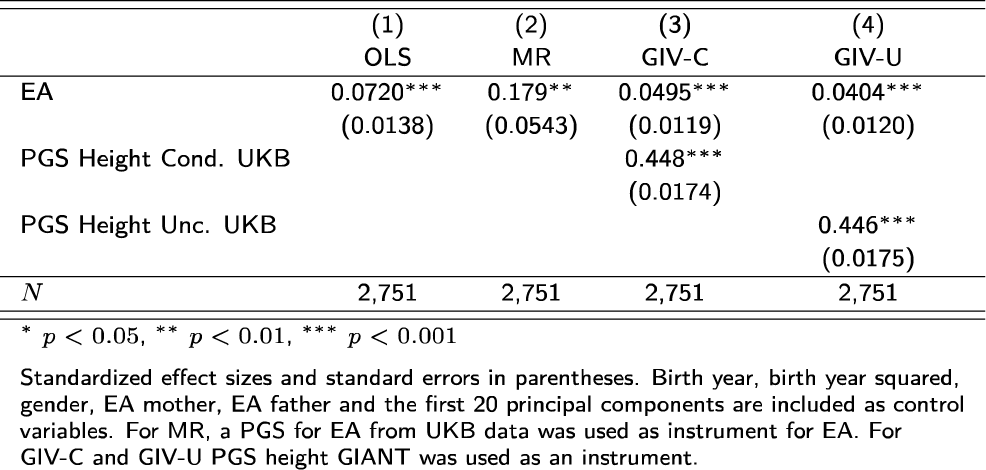
Estimates of the Effect of EA on Height.

These results also point towards a productive strategy for learning more about the true effect of height on EA. The demonstration that pleiotropy as well as environmental bias is affecting the estimates in Table 3 imply that there is no effective fix for MR; its genetic instruments are contaminated by pleiotropy and therefore cannot be used to adjust for environmental endogeneity. A convincing natural experiment that could randomize height within meaningful ranges would effectively address the question. Data on monozygotic twins would effectively control for pleiotropy and would be a productive estimation strategy assuming that there were not environmental factors that were both affecting height and EA and that would therefore bias the estimates from MZ twin data. Absent the existence of such data, the most productive strategy is arguably either to identify the relevant environmental factors to control or to search for data on siblings that could provide a more effective control of environmental variables via family fixed-effects while using GIV-U and GIV-C as proxy strategies to address bias from pleiotropy. With such data, the estimates from GIV-C and GIV-U would provide approximate bounds so long as the GIV-U estimate of the effect of EA on height was close to zero. If the negative control regression suggests bias from environmental sources, the GIV-U estimate will be more conservative than all the other estimators considered in this paper, and it will underpredict the true effect unless positive bias from both pleiotropy and from genetic-unrelated endogeneity is quite strong.

These results do not provide as clean and neat a conclusion as might be desired, though uncertainty is inevitable in the absence of experimental data or a valid IV. At the same time, the empirical example provides considerable insight into the implications of the available estimates. Our results strongly imply that OLS is providing an upwardly biased estimate of the effect of height on EA. They also strongly imply that the MR estimate suffers from pleiotropy bias and that MR is not an effective strategy for determining whether and to what extent height affects EA. Other studies suggest that pleiotropy between EA and height is not extremely high [18]. Our simulation results therefore suggest that GIV-U is either a plausible estimate for the effect of height on EA if genetic-unrelated endogeneity is relatively strong, or that it under predicts this effect if the endogeneity is weaker. If the estimate of GIV-U continued to be positive and statistically significant in a sibling fixed effects analysis, we would be rather confident that the effect is real and not an artifact of bias either from pleiotropy or from genetic-unrelated endogeneity. In other words, these results represent progress towards the goal of understanding how large is the social advantage provided by height in the process of EA given the existence of genetic confounds.^14^

## Conclusion

Accurate estimation of causal relationships with observational data is one of the biggest and most important challenges in epidemiology and the social sciences - two fields of inquiry where many questions of interest cannot be adequately addressed with properly designed experiments due to practical or ethical constraints. One important confound in non-experimental data comes from direct pleiotropic effects of genes on the exposure and the outcome of interest. Both OLS and MR yield biased results in this case. We proposed GIV regression as an empirical strategy that controls for such pleiotropic effects using polygenic scores. GIV regression uses standard IV estimation algorithms such as two-stage least squares that are widely available in existing statistical software packages. Our approach provides reasonable upper and lower bounds of causal effects in situations when pleiotropic effects of genes are the only source of bias. We showed that OLS, MR, and GIV regression yield biased estimates if both genetic and environmental sources of endogeneity are present. However, GIV regression still outperforms OLS and MR in this scenario. Furthermore, GIV regression can (and should) be combined with additional strategies that allow controlling for bias from purely environmental or behavioral factors, such as using co-variates or family-fixed effects. Together, these approaches can provide reasonable estimates of causal effects across a broad range of scenarios.

GIV regression is called for whenever an experimental design, a valid IV strategy, or a large-enough sample of MZ twins is not available and when pleiotropy is a potential problem - a situation that is frequently encountered in practice. The main requirements for GIV regression are a prediction sample that has been comprehensively genotyped and large-scale GWAS results for the outcome of interest from two non-overlapping samples. Thanks to rapidly falling genotyping costs that enable a growing availability of genetic data and large GWAS samples for many traits, these requirements become increasingly feasible for many applications. Indeed, the combination of new estimation tools and continued rapid advancements in genetics should provide a significant improvement in our understanding of the effects of behavioral and environmental variables on important socioeconomic and medical outcomes.

## ACKNOWLEDGMENTS

This research was facilitated by the Social Science Genetic Association Consortium (SSGAC) and by the research group on genetic and social causes of life chances at the Zentrum fur interdisciplinare Forschung (ZiF) Bielefeld. Data analyses make use of the UK Biobank resource under application’number 11425. We acknowledge data access from the Genetic Investigation of ANthropometric Traits Consortium (GIANT). We used data from the Health and Retirement Study (HRS), which is supported by the National Institute on Aging (NIA U01AG009740, RC2 AG036495, RC4 AG039029). HRS genotype data can be accessed via the database of Genotypes and Phenotypes (dbGaP, accession number phs000428.vl.pl). Researchers who wish to link genetic data with other HRS measures that are not in dbGaP, such as educational attainment, must apply for access from HRS. We are very grateful to Richard Karlsson Linner for help with the GWAS analyses in the UK Biobank, to Aysu Okbay for providing us with subsets of the GWAS meta-analysis on educational attainment, and to S. Fleur Meddens for constructing genetic principal components in the HRS data. We thank Daniel J. Benjamin, Jonathan P. Beauchamp, Benjamin Domingue, Lisbeth Trille Loft, Niels Rietveld, Eric Slob, Patrick Turley, Hans van Kippersluis, and Tian Zheng for productive discussions and comments on earlier versions of the manuscript. Furthermore, we thank Elliot Tucker-Drob for pointing us to the necessary correction of the heritability estimate in our model. The study was supported by funding from an ERC Consolidator Grant (647648 EdGe, Philipp D. Koellinger).

## Footnotes

1 Two other conditions that valid IVs need to satisfy are monotonicity (everyone who is affected by the IV is affected in the same direction) and the stable unit treatment value assumption (SUTVA): the “treatment” of one unit does not affect the outcome variable for other units.

2 Note that a clean experimental design which randomizes people into groups based on body height or EA is not possible. Thus, any attempt to study the causal relationship between the two variables must rely on observational data and naturally occurring experiments like the genetic endowment of individuals, which we exploit here.

3 Unfortunately, that is the reality in most social scientific and epidemiological studies that use non-experimental data.

4 i.e., the proportion of variance in a trait that is due to linear effects of currently measurable SNPs.

5 For an alternative approach to correcting attenuation bias based on the use of multiple indicators in a structural equation modeling framework, see [61].

6 We drop this assumption later. Also, we drop the subscript on the coefficients for the exogenous control variables *X* below when it would not lead to confusion.

7 Classic MR typically does not use PGS as instruments. Instead, the idea is to use single genetic variants that are known to affect the exposure via well-understood biological mechanisms that make it unlikely to violate the exclusion restriction. In practice, limited knowledge about the biological function of most genes make it difficult to argue that direct pleiotropic effects of the gene on the exposure and the outcome of interest exist.

8 Fixed effects estimation with panel data would also preclude MR type strategies because the IV does not vary over time, and genetic indicators for *T* would generally have a weak relationship to changes in *T* over time. Fixed effects regressions based on other strategies (e.g., sibling or neighborhood fixed effects models) would not control for pleiotropy. We discuss these strategies at greater length below.

9 An earlier version of our paper pursued this approach and called it GIV regression. However, we later found that controlling for *T* in a GWAS for *y* induces a correlation of *S*_*y*1|*XT*_ with *S*_*T*_ that invalidates the latter as an IV. The new version of GIV-U and GIV-C regression we describe below does not have this problem because it does not use a genetic instrument for *T* anymore. Instead, GIV-U and GIV-C both rely on a proxy-control strategy that only uses an instrument for *S*_*y*1|*XT*_ or *S*_*y*1|*X*_ to correct for measurement error in these proxies for 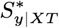 and 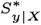.

10 It is possible to have finite sample bias that disappears asymptotically, in which case the estimator is consistent. We use the expectation formula instead because it is arguably more straightforward to understand.

11 The computer code for these simulations is available at https://github.com/cburik/GIVsim.

12 GREML yields unbiased estimates of SNP-based heritability that are not affected by attenuation, see [65].

13 Results from [18] and [15] suggest a genetic correlation between height and EA of about 0.15.

14 Obviously, none of the estimation strategies we discussed hens address possible bias from non-random selection into samples.

